# Partial information transfer from peripheral visual streams to foveal visual streams is mediated through local primary visual circuits

**DOI:** 10.1101/2023.10.12.561237

**Authors:** Andrea I. Costantino, Benjamin O. Turner, Mark A. Williams, Matthew J. Crossley

## Abstract

A classic view holds that visual object recognition is driven through the *what* pathway in which perceptual features of increasing abstractness are computed in a sequence of different visual cortical regions. The cortical origin of this pathway, the primary visual cortex (V1), has a retinotopic organization such that neurons have receptive fields tuned to specific regions of the visual field. That is, a neuron that responds to a stimulus in the center of the visual field will not respond to a stimulus in the periphery of the visual field, and vice versa. However, despite this fundamental design feature, the overall processing of stimuli in the periphery – while clearly dependent on processing by neurons in the peripheral regions of V1 – can be clearly altered by the processing of neurons in the fovea region of V1. For instance, it has been shown that task-relevant, non-retinotopic feedback information about peripherally presented stimuli can be decoded in the unstimulated foveal cortex, and that the disruption of this feedback – through Transcranial Magnetic Stimulation or behavioral masking paradigms – has detrimental effects on same/different discrimination behavior. Here, we used fMRI multivariate decoding techniques and functional connectivity analyses to assess the nature of the information that is encoded in the periphery-to-fovea feedback projection and to gain insight into how it may be anatomically implemented. Participants performed a same/different discrimination task on images of real-world stimuli (motorbikes, cars, female and male faces) displayed peripherally. We were able to decode only a subset of these categories from the activity measured in peripheral V1, and a further reduced subset from the activity measured in foveal V1, indicating that the feedback from periphery to fovea may be subject to information loss. Functional connectivity analysis revealed that foveal V1 was functionally connected only to the peripheral V1 and not to later-stage visual areas, indicating that the feedback from peripheral to foveal V1 is likely implemented by neural circuits local to V1.

## 1. Introduction

The ability to quickly and accurately recognize objects in the periphery of the visual field is of fundamental importance in recognizing and evading threats. Despite the importance of this ability, peripheral vision provides an impoverished representation of visual information relative to the fovea [1, 2, 3, 4, 5]. How might the visual system cope with such impoverished information in order to accomplish fast and accurate peripheral object recognition?

Recent work suggests that the answer to this question may lie in feed-back projections from peripheral to foveal visual areas. In a seminal study, [6] found that foveal cortex contained non-retinotopically mapped information about the shape of visual stimuli presented in the periphery, and that the magnitude of this information was correlated with visual discrimination performance. Subsequent studies established a causal link between the presence of this information and discrimination performance. First, high intensity rTMS over foveal cortex [7] impairs shape discrimination in the periphery. Second, an image presented centrally (to activate foveal cortex) after stimulus onset can impair performance if it is different from the images presented in the periphery, and improve performance if it is the same [8, 9, 10, 11]. Importantly, both the TMS and the behavioral manipulation only have an effect when applied in a specific time window (e.g., 350 ms after stimulus presentation in TMS, and 100-150 ms after stimulus presentation in the behavioral studies). The timing of both these effects is consistent with the idea that information arriving in the periphery is subsequently fed to foveal cortex along a multi-synaptic pathway.

However, the precise anatomical and functional nature of this feedback mechanism remains largely mysterious. Is it implemented as lateral projections between neurons entirely within primary visual cortex (V1), or is it instead implemented by projections from higher-order areas back to lower-order areas? In the former case we would expect activity in the fovea to encode only perceptual features (e.g., object shape and color), whereas in the latter case we would expect it to encode higher level category information (e.g., whether a viewed object is a face or a car). Existing studies are not well suited to answer these questions because as stimuli they have all used novel abstract visual objects for which there is no intrinsic higher-level category structure [12], leaving only perceptual differences between stimuli to drive neural and behavioral output.

Here, we examined peripheral object discrimination using stimuli that were composed of familiar objects (male faces, female faces, cars, and motorbikes) that have a clear hierarchical categorical structure. Male faces and female faces belong to the superordinate category ‘faces’ while cars and motorbike belong to the superordinate category ‘vehicles’. From a categorical perspective, the distinction between vehicles and faces is likely quite pronounced. However, the distinction between male and female faces, as well as between cars and motorbikes, might be comparatively less pronounced. These objects also display systematic variations in perceptual similarity, which differ from their categorical relationships. Specifically, male and female faces exhibit a high degree of perceptual similarity, cars and motorbikes are moderately similar, and faces and vehicles are the least similar.

Using fMRI and multivariate decoding techniques, we attempted to decode the category of these stimuli from key regions of interest. In peripheral V1, we expected to most successfully decode those stimuli drawn from categories that are very perceptually distinct. On the other hand, in higher-order visual areas (e.g., within the inferior temporal cortex) we expected to most successfully decode stimuli that are most categorically distinct. We also performed a functional connectivity analysis to identify whether the foveal region of V1 was functionally connected to peripheral V1 or to later-stage visual areas.

If feedback is implemented as lateral projections within V1, then we expect to find similar decoding abilities in foveal V1 as we find in peripheral V1. Similarly, if feedback is from higher-order peripheral areas, then we expect to find similar decoding ability in foveal V1 as we find in these higher-order areas. However, it is possible that – regardless of whether sourced from early or late visual areas – feedback from peripheral to foveal processing streams is only of partial representations (i.e., it does not preserve all available information). In this case, the decoding results obtained from fovea V1 may not look exactly like either aforementioned ROI.

Our results demonstrate that (1) feedback projections from peripheral V1 to foveal V1 are operative with real-world stimuli, (2) the pattern of decoding likely reflects the dominance of perceptual similarity in the ROIs and circuits we examined, (3) fovea V1 is functionally connected to peripheral V1 but not later-stage visual areas, and (4) the information that is relayed from peripheral stream to fovea V1 is only partial. These results are most consistent with the idea that feedback from periphery to fovea may be implemented within local V1 circuits (but see “Limitations and future directions” in the discussion). We discuss these findings in light of current models of visual processing, and speculate how these models might be amended to accommodate this new data.

## 2. Methods

### 2.1. Participants

Twenty-four participants were recruited from Macquarie University for this study. Participants were fluent in English, näıve to the purpose of the study and had normal or corrected-to-normal vision. Following Macquarie University Hospital standard procedure, all were screened for any fMRI contraindication and history of neurological impairment before being scanned. Participants had no metal permanently fixed to their body (e.g. dental work not including fillings such as braces, metal plates or screws) and were free of history involving neurological insult. The Macquarie Human Research Ethics Committee (Medical Sciences) approved this study, and each participant was asked to read and sign an informed consent form. Participants received $20 per hour as reimbursement for their participation in the study.

Stimuli were images of male faces, female faces, motorbikes, and cars. We adopt a nomenclature in which a *category-level* distinction between stimuli is given by whether or not the stimulus is a face or a vehicle, and a *subcategory* distinction is given by whether the face is male or female or whether the vehicle is a car or a motorbike (Fig. 1C). These stimuli were chosen in order to disentangle low-level perceptual information from high-level category information (more on this in section 2.7).

**Figure 1:**
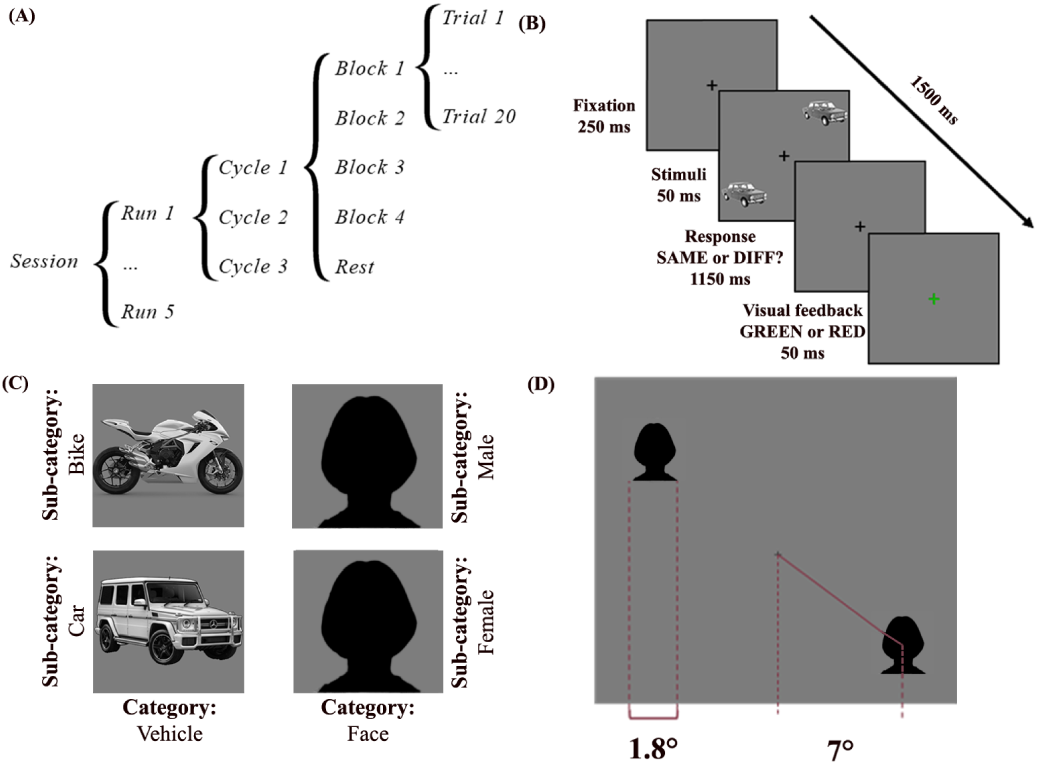
Faces are obscured due to the journal policy. (A) Each subject performed one scanning session consisting of 5 runs, 3 cycles per run, 5 blocks per cycle, and 20 trials per block. See text for further details. (B) An example trial from the experimental task showing a block in which the category was ‘vehicles’, the sub-category was ‘car’, and the trial type was ‘same’. (C) Each stimulus is part of a sub-category (i.e., bike, car, female or male), and each sub-category belongs to one of two categories (vehicle, face). (D) Stimuli were presented at 7 visual degrees eccentricity and measured 1.8 visual degrees on average.

60 Caucasian face images (30 male, 30 female) with neutral facial expressions were randomly selected from The Karolinska Directed Emotional Faces database [13]. 60 Vehicles (30 cars, 30 motorbikes) images with similar orientation and point of view were downloaded from online repositories. All images were resized to 700 *×* 700 px, converted to gray scale and equated in brightness and contrast using custom scripts written in MATLAB [14].

The stimuli were presented on a Cambridge Research System BOLD-Screen 32 LCD for fMRI (31.55 inch diagonal, 1920 *×* 1080 pixels, RGB color, 120Hz frame rate). The screen was placed in the MRI room and the participant could see the screen through a mirror placed inside the fMRI scanner (distance from eye to screen via mirror: 1120 mm). Each stimulus’ perceived size was *∼* 1.8° (visual degrees), presented at 7° diagonal eccentricity. These parameters matched those used in previous studies [7, 10, 6].

An Eyelink 1000 Remote Eye-tracker (long-range) was used to monitor participants’ eye movements (sampling rate: 1000 Hz; eye recorder: right; 5-points calibration).

### 2.2. Procedure

Participants were asked to attend two separate sessions for this study. In the first session, participants were familiarized with the task and eyetracking procedure. Participants performed two practice runs (see below for a description of a run) in the computer lab at the Australian Hearing Hub. The second session was run at the fMRI facility of the Macquarie Medical Imaging center. During this session, participants performed two tasks while lying in the scanner: the localizer task and the experimental task.

#### 2.2.1. Localizer task

Before the experimental task, participants performed a 1-back sequential matching task. In each trial, a single stimulus was presented at the center of the screen for 500 ms, followed by a 500 ms response window. Participants were asked to press a button with the index finger during the response window if the same stimulus was presented two consecutive times.

Stimuli were drawn from the same categories and sub-categories as those used for the experiment ask, but stimuli used for the localizer task were not used for the experiment task. As in previous studies [6], the average stimulus size was 8 visual degrees.

Participants were asked to perform one session including ten cycles, where each cycle included two blocks (one for each category) and one rest block. The total length of the task was 8 minutes, and each block lasted 20 seconds. The functional data obtained from the localizer task was then processed and combined with functional parcels in order to identify two distinct Regions of Interest (ROIs): a face-selective ROI (Fusiform Face Area or FFA), and an object-selective ROI (Lateral Occipital Complex or LOC). See section 2.5 for more details.

#### 2.2.2. Experiment task

After completing the localizer task, participants were asked to complete the experimental task. Stimuli in this task were pairs of images from the same sub-category displayed simultaneously either on the top-right and bottom-left corners (for participants assigned to the ‘normal’ orientation condition) or on the top-left and bottom-right corners of the screen (for participants assigned to the ‘inverted’ orientation condition). The experiment task was performed over 5 functional MRI image acquisition runs, 3 cycles per run, 5 blocks per cycle, and 20 trials per block (Fig. 1A).

On every trial of the experimental task, participants were asked to decide whether two images that were briefly and simultaneously displayed in the periphery of the visual field were the same or different (Fig. 1D). Trials consisted of a 250 ms fixation period, 50 ms of stimulus presentation, a 1150 ms response window, and 50 ms of feedback presentation (Fig. 1B). Thus, each trial was exactly 1500 ms in duration. A 0.5° fixation cross centered on the screen was displayed throughout the entire duration of each trial, including during stimulus presentation and throughout the response window, and participants were instructed to maintain their gaze on this fixation cross at all times. During the fixation period, only the fixation cross was visible. Following stimulus offset, participants were required to make a same/different judgment within the response window. Responses were made by pressing one button with their index finger for ‘same’ or a different button with their middle finger for ‘different’. Responses made outside the response window were recorded as incorrect responses. During feedback presentation, the fixation cross turned green for correct responses or red for incorrect responses and remained colored. After completing a cycle (see below), participants were required to press a button to begin the next.

‘Same’ trials were constructed by pairing two identical stimuli belonging to the same sub-category, while ‘different’ trials were constructed by pairing two different randomly selected stimuli belonging to the same sub-category (i.e., within-subcategory discrimination).

The 20 trials constituting a block were 10 ‘same’ trial types and 10 ‘different’ trial types, and the order in which these stimuli were presented was randomized within each block. The 5 blocks per cycle were composed of 4 *×* 30 sec experimental blocks (one for each sub-category) and 1 *×* 20 sec rest block.

Only one sub-category of stimuli was presented in each block. For each subject, the order in which sub-categories were presented was randomly sampled without replacement from the set of all possible orders that span the entire experimental task. This set is made up of 60 possible sequences (5 runs *×* 3 cycles *×* 4 experiment blocks *×* 1 sub-category per block). Note that since our number of participants (*N* = 25) is less than this set size, every participant received a unique experiment-wide sub-category order.

There were 240 trials per run (3 cycles per run *×*4 blocks per cycle *×* 20 trials per block). The same set of stimuli – and therefore the same set of trial types – was presented to each subject in every run. Since only one sub-category of stimuli was presented in each block, 60 unique trials were shown for each sub-category during a run.

### 2.3. fMRI parameters

fMRI scans were acquired on a 3T Siemens Verio MRI scanner at Macquarie Medical Imaging (MMI), Macquarie University Private Hospital, Sydney, using a 32-channel head coil. Functional scans used a sequential descending T2*-weighted echo-planar imaging (EPI) acquisition sequence: acquisition time 2000ms; interleaved (Multiband acquisition, 3 slices); echo time 30.8ms; resolution 1.9*×*1.9*×*1.9mm + 20% spacing; flip angle 84 deg; 51 slices. The high-resolution structural image was T1-weighted MPRAGE; slice thickness 1mm, resolution 1*×*1mm, flip angle: 9 deg; echo time: 3.03ms; TR: 1800ms.

Once participants entered the room where the fMRI scanner is located, they were instructed to lie down on a bed and were given earplugs to reduce the noise inside the scanner. Padding foam was placed around the partici pant’s head to ensure a stable head position, and participants were shown a safety button to press to interrupt scanning at any time. Before the actual testing session started, a 1mm isotropic voxel structural scan was acquired. After the structural MRI scanning, participants performed the experimental task. The total scanning time was *∼*50 min (5 min for the T1 weighted structural + 45 min for the T2* functional). Participants had a one-minute break between each run.

### 2.4. fMRI QA and preprocessing

*HeuDiConv* [15] was used to convert the raw images from the scanner into a Brain Imaging Data Structure (BIDS) compliant database [16]. Image Quality Assessment (QA) was performed using *MRIQC* [17], and *MRIQ-Ception* [18] was used to compare Image Quality Metrics (IQM) against normative data. After visual inspection of the structural and functional IQM distributions, no subjects were removed from further analysis.

Results included in this manuscript come from the preprocessing steps described in the following two subsections all of which were performed using *fMRIPrep* 20.2.1 [19, 20] – RRID:SCR 016216 – which is based on *Nipype* 1.5.1 [21, 22] – RRID:SCR 002502. Many internal operations of *fMRIPrep*, mostly within the functional processing workflow, use *Nilearn* 0.6.2 [23], RRID:SCR 001362 . For more details of the pipeline, see the section cor-responding to workflows in *fMRIPrep*’s documentation.

#### 2.4.1. Anatomical data preprocessing

A total of 1 T1-weighted (T1w) images were found within the input BIDS dataset for each subject. The T1-weighted (T1w) image was corrected for intensity non-uniformity (INU) with `N4BiasFieldCorrection` [24], distributed with ANTs 2.3.3 [25] – RRID:SCR 004757 – and used as T1w-reference throughout the workflow. The T1w-reference was then skull-stripped with a *Nipype* implementation of the `antsBrainExtraction.sh` workflow (from ANTs), using OASIS30ANTs as target template. Brain tissue segmentation of cerebrospinal fluid (CSF), white-matter (WM) and gray-matter (GM) was performed on the brain-extracted T1w using fast [26] – FSL 5.0.9, RRID:SCR 002823. Brain surfaces were reconstructed using recon-all [27] – FreeSurfer 6.0.1, RRID:SCR 001847 – and the brain mask estimated previously was refined with a custom variation of the method to reconcile ANTs-derived and FreeSurfer-derived segmentations of the cortical gray-matter of Mindboggle [28] – RRID:SCR 002438. Volume-based spatial normalization to one standard space (MNI152NLin2009cAsym) was performed through nonlinear registration with antsRegistration (ANTs 2.3.3), using brain-extracted versions of both T1w reference and the T1w template. The following template was selected for spatial normalization: *ICBM 152 Nonlinear Asymmetrical template version 2009c* [29] – RRID:SCR 008796, Template-Flow ID: MNI152NLin2009cAsym).

#### 2.4.2. Functional data preprocessing

For each of the 6 BOLD runs found per subject (across all tasks and sessions), the following preprocessing was performed. First, a reference volume and its skull-stripped version were generated using a custom methodology of *fMRIPrep*. Susceptibility distortion correction (SDC) was omitted. The BOLD reference was then co-registered to the T1w reference using bbregister (FreeSurfer) which implements boundary-based registration [30]. Co-registration was configured with six degrees of freedom. Head-motion parameters with respect to the BOLD reference (transformation matrices, and six corresponding rotation and translation parameters) are estimated before any spatiotemporal filtering using `mcflirt` [31] – FSL 5.0.9. BOLD runs were slice-time corrected using `3dTshift` from AFNI 20160207 [32] – RRID:SCR 005927. The BOLD time-series (including slice-timing correction when applied) were resampled onto their original native space by applying the transforms to correct for head-motion. These resampled BOLD time-series will be referred to as *preprocessed BOLD in original space*, or just *preprocessed BOLD*. The BOLD time-series were resampled into standard space, generating a *preprocessed BOLD run in MNI152NLin2009cAsym space*. First, a reference volume and its skull-stripped version were generated using a custom methodology of *fMRIPrep*. Several confounding time-series were calculated based on the *preprocessed BOLD* : framewise displacement (FD), DVARS and three region-wise global signals. FD was computed using two formulations following Power – absolute sum of relative motions [33] – and Jenkinson – relative root mean square displacement between affines [31]. FD and DVARS are calculated for each functional run, both using their implementations in *Nipype* following the definition by [33]. The three global signals are extracted within the CSF, the WM, and the whole-brain masks. Additionally, a set of physiological regressors were extracted to allow for component-based noise correction (*CompCor*, [34]). Principal components are estimated after high-pass filtering the *preprocessed BOLD* time-series (using a discrete cosine filter with 128s cut-off) for the two *CompCor* variants: temporal (tCompCor) and anatomical (aCompCor). tCompCor components are then calculated from the top 2% variable voxels within the brain mask. For aCompCor, three probabilistic masks (CSF, WM and combined CSF+WM) are generated in anatomical space. The implementation differs from that of Behzadi et al.*∼* in that instead of eroding the masks by 2 pixels on BOLD space, the aCompCor masks are subtracted a mask of pixels that likely contain a volume fraction of GM. This mask is obtained by dilating a GM mask extracted from the FreeSurfer’s *aseg* segmentation, and it ensures components are not extracted from voxels containing a minimal fraction of GM. Finally, these masks are resampled into BOLD space and binarized by thresholding at 0.99 (as in the original implementation). Components are also calculated separately within the WM and CSF masks. For each CompCor decomposition, the *k* components with the largest singular values are retained, such that the retained components’ time series are sufficient to explain 50 percent of variance across the nuisance mask (CSF, WM, combined, or temporal). The remaining components are dropped from consideration. The head-motion estimates calculated in the correction step were also placed within the corresponding confounds file. The confound time series derived from head motion estimates and global signals were expanded with the inclusion of temporal derivatives and quadratic terms for each [35]. Frames that exceeded a threshold of 0.5 mm FD or 1.5 standardized DVARS were annotated as motion outliers. All resamplings can be performed with *a single interpolation step* by composing all the pertinent transformations (i.e., head-motion transform matrices, susceptibility distortion correction when available, and co-registrations to anatomical and output spaces). Gridded (volumetric) resamplings were performed using `antsApplyTransforms` (ANTs), configured with Lanczos interpolation to minimise the smoothing effects of other kernels [36]. Non-gridded (surface) resamplings were performed using mri vol2surf (FreeSurfer).

**Figure 2:**
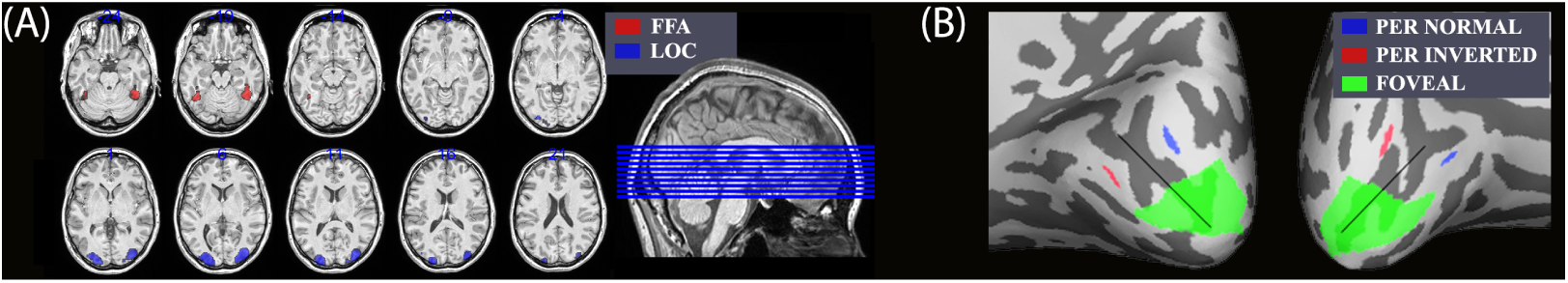
ROIs plotted on individual subject anatomical scan. (A) Functional ROIs. To generate individuals ROIs, parcels from the GCSS analysis were used as masking images on the statistical (t) activation maps resulting from the relevant contrasts in the SPM first level analysis (Red: ‘FFA’; Blue: ‘LOC’). (B) Retinotopic ROIs. (Green: ‘Foveal’, 4°; Blue: ‘Peripheral normal’; Red: ‘Peripheral inverted’; Black line: Calcarine Sulcus)

### 2.5. Regions of interest (ROIs)

We defined 5 ROIs – FFA, LOC, peripheral V1 (corresponding to stimulus location), peripheral V1 (corresponding to where stimuli were not presented), and foveal V1 – within which we restricted our multivariate decoding analyses. FFA and LOC were defined on the basis of functional runs, and the V1 ROIs were defined retinotopically. See Fig. 2 for an illustration of the ROIs.

#### 2.5.1. Functional ROIs

To identify the functional category-selective ROIs, participants performed a localizer task (see section 2.2 for details). To algorithmically define the ROIs for each subject, a set of ‘parcels’ (broad regions involved in faces and objects processing) were obtained from the group data using a group-constrained subject-specific (GCSS) analysis. The GCSS was carried out by adapting an algorithm already used in [37], and the procedure has been extensively described in [38]. To identify the parcels of interest, a first level analysis (GLM) was performed in SPM12 on the smoothed data (FWHM = 5mm) of the localizer task. A set of confounds – specifically, ’*rot_x_*’, ’*rot_y_*’, ’*rot_z_*’, ’*trans_x_*’, ’*trans_y_*’, ’*trans_z_*’ – obtained from *fmriprep* were included in the model to remove the variance explained by motion confounds. The relevant contrasts (Faces ¿ Vehicles, Vehicles ¿ Faces) were generated in SPM for each participant, and a statistical *t* map was generated for each contrast / subject. The *t* maps were converted to *z* maps in MATLAB. Each statistical map was then thresholded (at *p* = 0.001 for Faces ¿ Vehicles, *p* = 0.000001 for Vehicles ¿ Faces) and binarized, and all subjects’ statistical maps were overlaid. The probability map obtained from the overlay was smoothed (FWHM = 6mm) and thresholded at two subjects. Finally, a watershed algorithm implemented in SPM-ss [39] was applied to generate parcels based on local maxima in the probability maps. The resulting parcels were considered representative, as the voxels were significant in at least 80% of the subjects and corresponded to the expected anatomical locations for left or right FFA, and left or right LOC. After generating the parcels, the single subjects’ ROIs were defined iteratively in Python 3.7.9 with custom scripts based on *Nibabel* [40] and *Nilearn* [23] by combining the statistical *t* maps generated by SPM thresholded at *p <* 0.01 (Faces ¿ Vehicles, Vehicles ¿ Faces) and the parcels generated by the GCSS analysis. Therefore, 4 ROIs were defined for each subject: *FFA_l_, FFA_r_, LOC_l_, LOC_r_*. Left and right masks were combined in a single bilateral mask for further analysis.

#### 2.5.2. Retinotopic ROIs

The retinotopic ROIs were generated using *Neuropythy* [41] and the ‘Benson-2014’ atlas [42] to predict brain activations in V1 corresponding to visual stimuli with given eccentricity and polar angle. The ROIs were defined on the subject cortical surface and then transformed to subject anatomical volume space and resampled to match the functional resolution. We defined 2 peripheral and 8 foveal ROIs.

The peripheral ROIs represented the predicted activation in V1 elicited by two disks (size: 2°) at 7° eccentricity from the fovea and +45°/+225° from the Upper Vertical Meridian (UVM); and two disks (size: 2°) at 7° eccentricity from the fovea and -45°/-225° from the UVM. The retinotopic location of these ROIs corresponded to the location of the stimuli presented during the experimental task in ‘normal’ and ‘inverted’ stimuli orientation, respectively. For simplicity, here we will refer to ‘Peripheral’ and ‘Opposite’ ROIs respectively to indicate the ROI representing the stimulus location and the ROI representing the opposite to the stimulus location (i.e., no visual stimulation).

Choosing an appropriate foveal ROI is a challenge because we don’t know the exact size of the foveal V1 region that the peripheral-stream to foveal-stream feedback projection targets. If we choose an ROI that is too small, it won’t capture all the relevant information and decoding will be poor. On the other hand, if we choose an ROI that is too large, it will include irrelevant information and decoding will also be poor ^1^. Therefore, we generated eight foveal ROIs ranging in size from 1 to 4 visual degrees (average number of voxels: 135.17, 300.42, 457.42, 603.46, 738.29, 857.67, 964.5, 1063.25) using the same method employed for the creation of peripheral ROIs. To find the most appropriately sized Foveal ROI, we fit a mixed effects second order polynomial regression to the decoding accuracies from each participant, taking decoding accuracy as the observed variable and ROI size as the predictor variable. This is captured by the following regression equation:

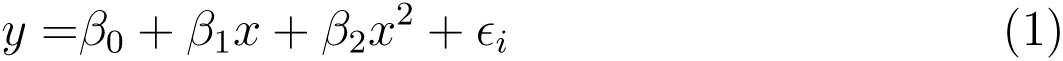

Here, *β*_0_, *β*_1_, and *β*_2_ are group-level fixed effect regression coefficients and *E_i_* is the random intercept for subject *i*. We took the Foveal ROI that corresponded to the peak predicted decoding accuracy as the most appropriate Foveal ROI for use in subsequent analyses. Based on this analysis, we focused on the 2° foveal ROI for subsequent analyses.

### 2.6. fMRI Multi-Voxel Pattern Analysis

We performed multi-voxel pattern analysis (MVPA) independently for each subject and for each ROI using ROI-masked beta-maps as multi-voxel input patterns. As input patterns for category-level classification (i.e., Faces vs. Vehicles), we used beta-maps obtained from a GLM analysis using *Face* and *V ehicle* as regressors, and *rot_x_*, *rot_y_*, *rot_z_*, *trans_x_*, *trans_y_*, *trans_z_* as confounds. As input patterns for sub-category-level classification (i.e., Male vs. Female vs. Car vs. Bike) and for within sub-category classification (i.e., Female vs. Male, Bike vs. Car), we used beta-maps obtained from a GLM using *Male*, *Female*, *Car*, and *Bike* as regressors, and *rot_x_*, *rot_y_*, *rot_z_*, *trans_x_*, *trans_y_*, *trans_z_* as confounds. Both GLM analyses were performed in SPM12. Events were pooled across all blocks and cycles such that we obtained one beta-map per run per sub-category. Finally, before submitting these beta maps to MVPA, from each value in the beta maps of the compared conditions, we subtracted the average value across all voxels in the corresponding maps (for details, see [43]) and NaNs were converted to zero.

Using sklearn [44], we trained linear Support Vector Machines (SVM) to classify multi-voxel activation pattern differences in the beta images described above. For every subject, the classifier accuracy was obtained for each ROI and contrast. The classifiers were trained on the data of 4 experimental runs and tested on the remaining run (leave-one-run-out) in each of the 5 folds. The average accuracy was then computed across these 5 folds to yield the final accuracy for that subject and classifier. A one-sample *t*-test was then performed on these average accuracies against chance level (25% or 50% depending on the condition) to investigate the classifier ability to discriminate between categories and sub-categories at the group level.

### 2.7. MVPA predicted from a model of early visual areas

We compared the results from the MVPA methods described above to those predicted from the first convolution layer in AlexNet [45] – a deep convolutional neural network – from the PyTorch [46] model zoo. The purpose of this comparison was to help formalise what pattern of decoding we would expect to get from a system operating only on perceptual features, which is the same pattern that we would expect in peripheral V1. Note that the the first convolution layer in AlexNet – like the primary visual cortex in humans – possesses receptive fields that encode simple perceptual features of stimuli (e.g., spatial frequency and orientation) as shown in the top panel of Fig. 3 and it is for this reason that we used AlexNet to estimate the perceptual similarity structure of our stimuli (bottom panel of Fig. 3).

### 2.8. Stimuli and apparatus

All original 120 centered, gray scale, 700 *×* 700 px images employed in this study (i.e., bikes, cars, female and male faces) were used in this analysis. Images were normalized in all the three RGB channels using the mean and standard deviation of the ImageNet [47] Database (mean = [0.485, 0.456, 0.406], sd = [0.229, 0.224, 0.225]), and resized to match the required AlexNet input size (224 *×* 224 pixels). For each normalized image, we obtained an image-evoked network activation pattern by extracting the output of the first convolutional layer, reducing its dimension to 100 *×* 1 by applying truncated singular value decomposition (SVD), and adding zero-mean Gaussian random noise (see below for details on the standard deviation of this noise). We then performed a classification analysis similar to that described in the previous two sub-sections but using network activation patterns as predictors instead of fMRI beta maps. Specifically, we used stratified K-fold cross validation to generate 20 train / test splits, and trained a linear SVM to predict image category and sub-category from image-evoked network activation patterns. We repeated this procedure 50 times, each with a different sample of Gaussian random noise and with different train / test splits.

**Figure 3:**
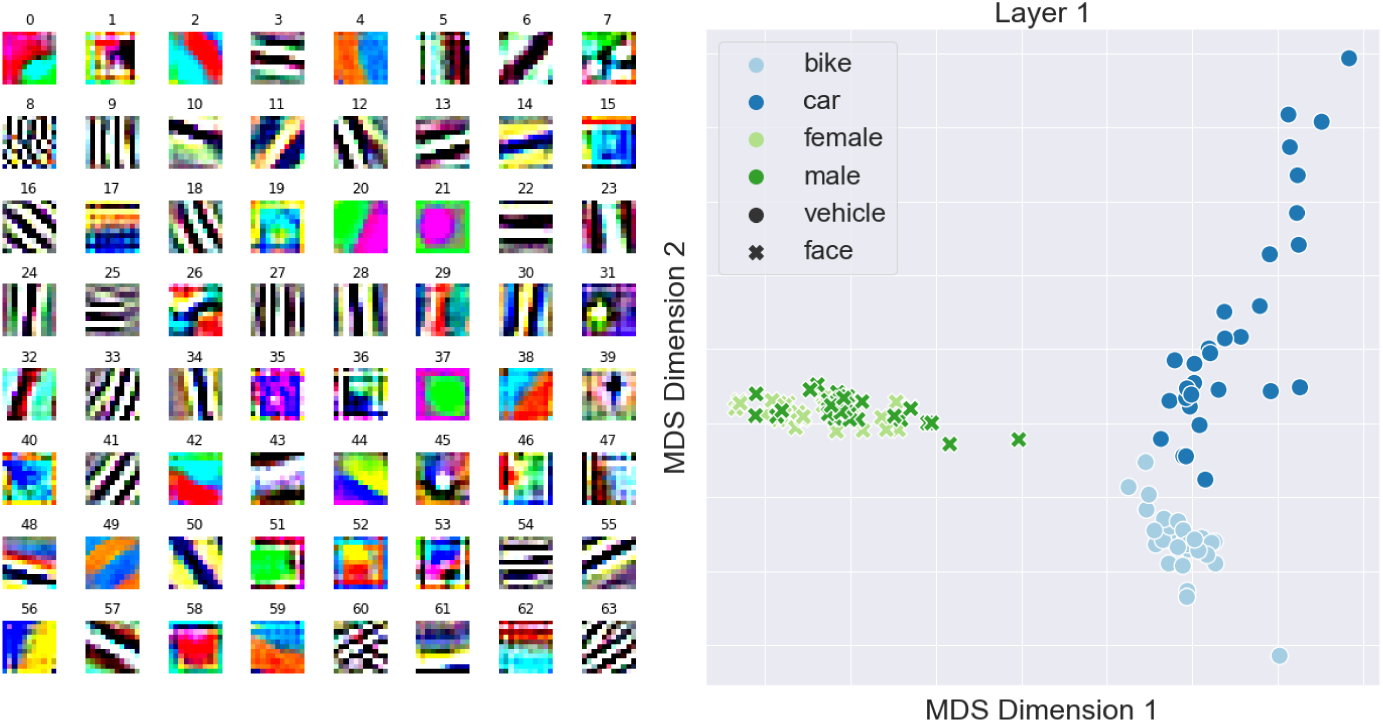
(Top) Weight matrices from the first convolution layer in AlexNet. Note that these weights appear very similar to a bank of gabor filters at different spatial frequencies and orientations, indicating that a image-induced activation of this layer in AlexNet will encode primarily perceptual features of the image. (Bottom) Perceptual similarity between all stimuli used in our experiment. See text for a detailed explanation of how activation patterns from AlexNet were converted into this 2-dimensional space.

The bar height in Fig. 4 are the average of all 20 *×* 50 classifiers, and the error bars are the standard deviation of all 20 *×* 50 classifiers divided by the square root of 20. Finally, we repeated this analysis for 5 different standard deviations of Gaussian random noise (*σ* = 0*, σ* = 50*, σ* = 100*, σ* = 150*, σ* = 200). The results in Fig. 4 correspond to *σ* = 100, which we chose to show because it led to classifier accuracy that was approximately midway between ceiling and floor. However, the results are qualitatively similar for every value of standard deviation explored (supplemental Fig. 1).

### 2.9. Functional connectivity via PPI

We performed a psychophysiological interaction analysis (PPI) to estimate functional connectivity between the fovea and each ROI (FFA, LOC, Opposite, and Peripheral). This analysis involves fitting a linear regression to predict the BOLD time series of the following form:

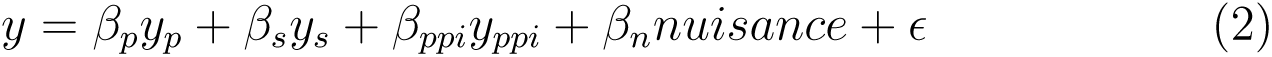

**Figure 4:**
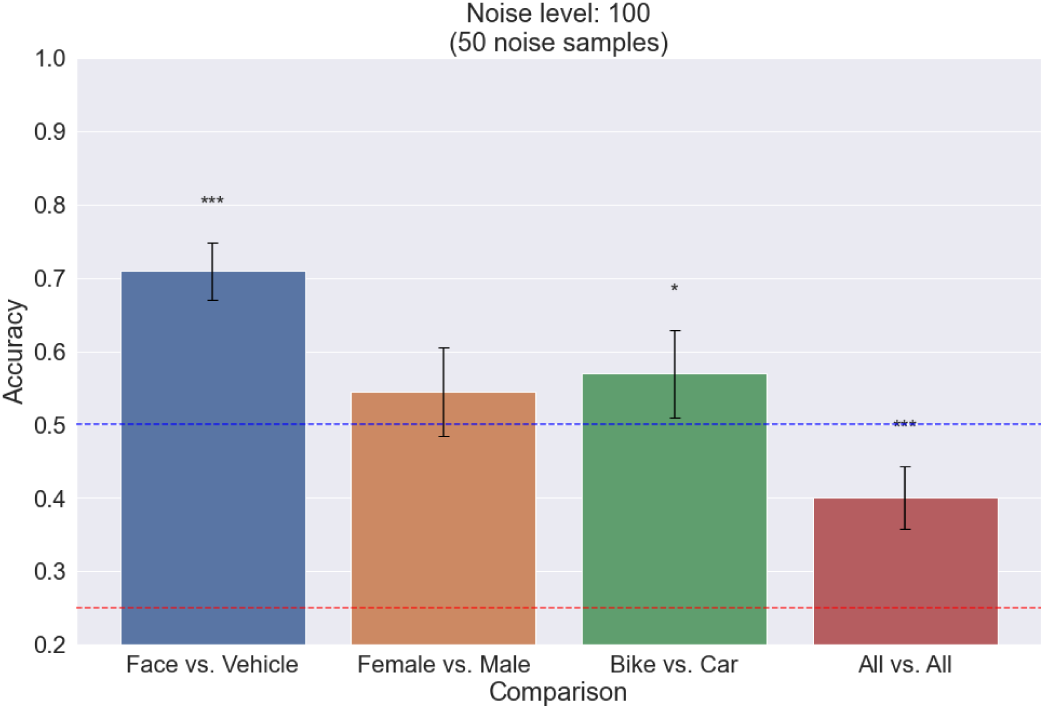
Decoder accuracy obtained from layer 1 of AlexNet for category level (Face vs. Vehicle, in blue), within-category faces (Male vs. Female, orange), within category vehicles (Car vs Bike, green), and sub-category level (Bike vs. Car vs. Female vs. Male, in red). Error bars: SEM as defined in the Perceptual model section. Blue line: category level chance (50%). Red line: sub-category level chance (25%). Asterisks (*) indicate statistical significance (e.g., * p < 0.05, ** p < 0.01, *** p < 0.001).

Here, the *y* term on the LHS is the BOLD time series from a target region (i.e., the foveal ROI). The first term on the RHS *y_p_*is a boxcar function convolved with a model HRF indicating when the stimulus is present and is referred to as the *psychological* term in a PPI analysis. We constructed this term such that the boxcar function takes the value 1 when any stimulus from any category or sub-category is displayed and 0 during the 20s rest periods. If the regression coefficient of *y_p_* is significantly greater than zero, then the analysis indicates that the target region shows task related activity, but says nothing of its connectivity with the seed region.

The second term on the RHS *y_s_*is the BOLD time series from a seed region (FFA, LOC, Opposite, or Peripheral) and is referred to as the *physiological* term in a PPI analysis. If the regression coefficient of *y_s_* is significantly greater than zero, then the analysis indicates that the target region correlates with the seed region, but still does not indicate functional connectivity because it cannot rule out the possibility that both regions are driven by a common third region.

The third term on the RHS *y_ppi_* is *y_s_* multiplied by a version of *y_p_* that is scaled and translated to span the interval [*−*1, 1]. That is,

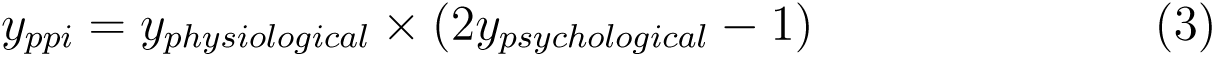

The term *y_ppi_* is referred to as the *psychophysiological interaction* term in a PPI analysis. If the regression coefficient of *y_ppi_* is significantly greater than zero, then the analysis indicates that the target region positively correlates with the seed region during task periods and not during rest periods (i.e., the target region shows context-specific correlation with the seed region). This is taken as evidence for functional connectivity.

Finally, *nuisance* regressors include head motion, drift, and run predictors and *E* is a normally distributed residual noise term. Overall, the coefficient of *y_ppi_*is the critical term to this analysis.

All BOLD time series were fully preprocessed as described in section 2.4 and further processed here by using PCA to reduce the dimension of each ROI time series from *n*_voxel_ *×n_T R_* to 1*×n_T R_*. The input data was centered but not scaled before being subjected to PCA and the resulting 1-dimensional component from each region was standardized to have unit variance.

This process yields a separate 1 *× n_T R_* signal for each ROI and for each subject. For each seed ROI (FFA, LOC, Opposite, and Peripheral), we estimated a linear mixed effects version of Eq. 2 in which each subject was given a random global intercept and random slope on each primary predictor variable (*y_p_*, *y_s_*, *y_ppi_*). That is, we estimated the following model:

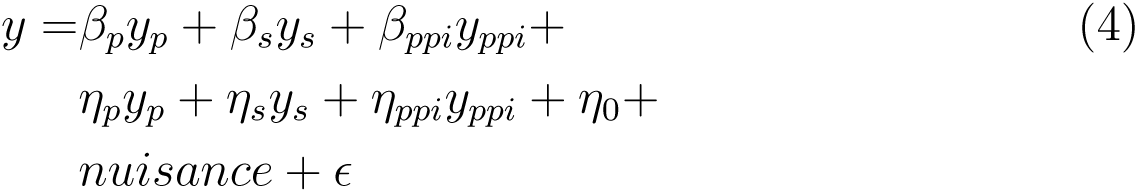

The terms in the first and last lines of Eq. 4 contain fixed effects terms and are identical to the corresponding terms in Eq. 2. The middle line of Eq. 4 contains random effects terms (i.e., per subject random slopes and per subject global intercept). Thus, all *β* terms are treated as constants to be estimated and all *η* terms are treated as normally distributed random variables with means and variances to be estimated. That is,

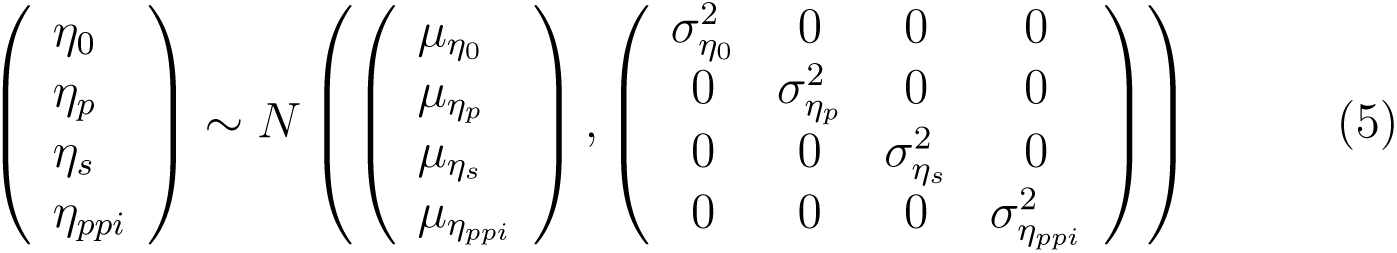

Note that each subject is modelled with their own random intercept and random slopes but that for this is not explicitly indicated in the nomenclature used in the above equations. In essence, every term on line 2 of Eq. 4 and every term anywhere in Eq. 5 should have a subscript indicating that there are as many of these expressions as there are subjects. We omitted this for simplicity.

The critical term of this analysis is again *β_ppi_*, the coefficient of the fixed effect *y_ppi_* regressor. We estimated the statistical significance of this term using two methods. First, we used Satterthwaite’s approximation to calculate degrees of freedom, t-statistics, and p-values for the fixed effect coefficients in Eq. 4 [48, 49].

Second, we used parametric bootstrap methods to compare the full Eq. 4 model to a reduced version in which the fixed effect *β_ppi_y_ppi_* was removed [50]. For this method, we generated 5000 samples of the likelihood ratio test statistic (LRT) and calculated p-values for rejecting the reduced model in favour of the full model as the fraction of simulated LRT-values that are larger or equal to the observed LRT value.

We also calculated p-values by (1) assuming that the LRT has a chi-square distribution, or (2) applying a Bartlett correction to the LRT from the mean of the simulated LRT-values, or (3) assuming that the LRT has a gamma distribution with mean and variance determined as the sample mean and sample variance of the simulated LRT-values, or (4) assuming that the LRT divided by the number of degrees of freedom has an F distribution where the denominator degrees of freedom are determined by matching the first moment of the reference distribution [50].

These methods for ascertaining the significance of the *β_ppi_* are appropriate because they produce acceptable Type I error rates (*α* = 0.05) even for smaller sample sizes [51, 52, 53].

### 2.10. Statistics and software

ANOVAs and *t*-tests were performed using Python 3.8 and *SciPy* [54]. The Python library *Scikit-learn* was used to perform PCA dimension reduction [55]. The R package *lme4* was used to implement linear mixed effect models [56], the *lmerTest* package was used to obtain p-values from these models [49], and the package *pbkrtest* was used to perform parametric bootstrap model comparison along with other LRT model comparison approaches [50].

## 3. Results

We collected Eye-tracking, fMRI and behavioral (same / different accuracy) data of each participant (*n* = 25) during the experiment task. We excluded one participant from further analysis due to fMRI technical issues. As the average fixation for the remaining (*n* = 24) participants was within 2 degrees radius from the fixation cross, we didn’t exclude any subjects based on Eye-tracking data (see supplemental Fig. 2).

### 3.1. Behavioral

The average behavioral accuracy across sub-categories was 67% (*n* = 24; *M_cars_*= .66*, M_bikes_* = .68*, M_male_* = .66*, M_female_* = .69). A one-way ANOVA on participants’ accuracy across sub-categories and runs revealed no significant difference in accuracy between sub-categories (*F* (3, 84) = .33*, p* = .81*, η*^2^ = 0.01) or runs (*F* (4, 105) = .83*, p* = .51*, η*^2^ = 0.03). In other words, stimuli from each subcategory are equally confusable. Note that, as described above, estimates for the activity per subcategory for the subsequent MVPA come from fitting a single regressor to each category or subcategory respectively. The results shown in Fig. 5 suggest that this averaging operation is equally valid across all subcategories. This in turn rules out as an alternative explanation for the MVPA results the possibility of differences in within-subcategory homogeneity, or equivalently, differences in the representativeness the within-subcategory average.

### 3.2. MVPA

Fig. 6 shows classifier accuracies in the main regions of interest (See also Table 1). In the FFA, SVM accuracy was significantly greater than chance when comparing all the sub-categories (Fig. 6 red bar: Bike vs. Car vs. Female vs. Male, *FFA* : *t*(23) = 10.3*, p < .*0001*, d* = 2.15) and categories (Fig. 6 blue bar: Faces vs. Vehicles, *FFA* : *t*(23) = 34.44*, p < .*0001*, d* = 7.18), but was not significantly greater than chance for within-category faces (Fig. 6 orange bar: Male vs. Female) or vehicles (Fig. 6 green bar: Car vs. Bike). These results indicate that between-category information (Faces vs. Vehicles) is encoded in FFA, which is consistent with its established role in face processing. Please see the “Neural architecture of periphery-to-fovea feedback” discussion section for comments on why we may have been unable to decode male faces from female faces in the FFA.

**Figure 5:**
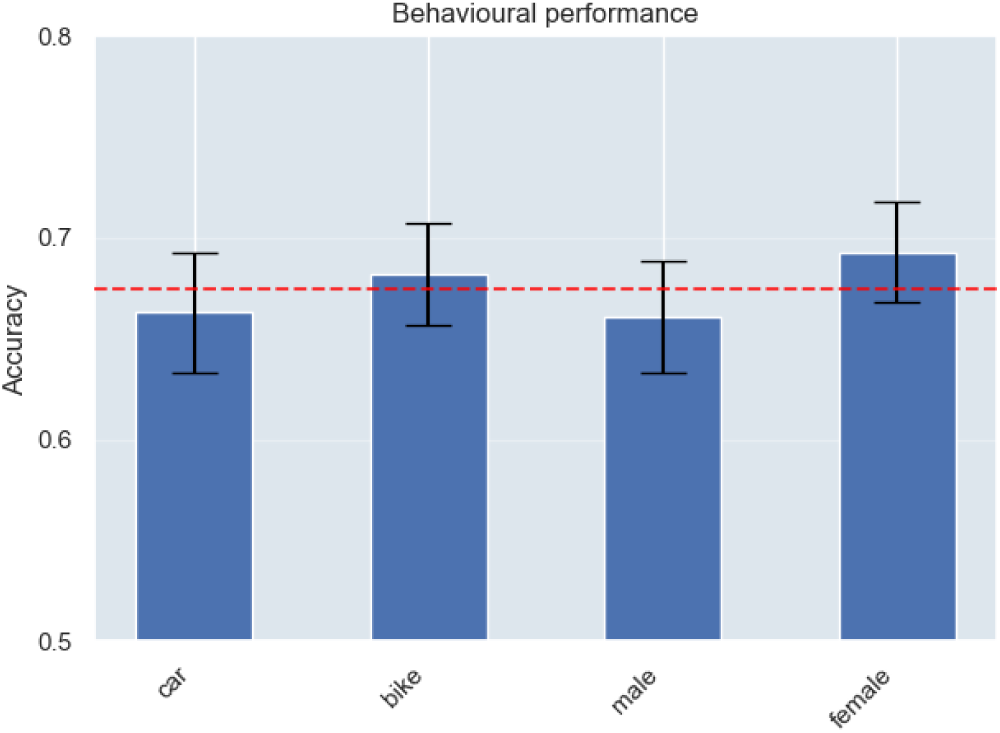
Average behavioral accuracy. Error bars: SEM. Red line: average accuracy across all categories.

In the LOC, SVM accuracy was significantly greater than chance when comparing all sub-categories (Fig.6 red bar: *LOC* : *t*(23) = 2.19*, p < .*05*, d* = 0.46), categories (Fig.6 blue bar: *LOC* : *t*(23) = 3.09*, p < .*01*, d* = 0.64), and for within-category vehicles (Fig.6 green bar: Car vs. Bike, *LOC* : *t*(23) = 2.22*, p < .*05*, d* = 0.46). SVM accuracy was not significantly greater than chance for within-category faces (Fig.6 orange bar: Male vs. Female). These results are consistent with previous results suggesting a central role of LOC in shape discrimination [57]. However, as noted below, this may also be driven by perceptual similarity more than by categorical encodings in LOC.

As expected, nothing was above chance in the Opposite ROI (i.e., opposite to the stimuli location) for any contrast. In the Peripheral ROI (i.e., stimuli location), we found greater than chance SVM accuracy when comparing categories (Fig.6 blue bar: ‘Face vs. Vehicle’ *PER* : *t*(23) = 2.02*, p < .*05*, d* = 0.42), all sub-categories (Fig.6 red bar: ‘Bike vs. Car vs. Female vs. Male’ *PER* : *t*(23) = 2.09*, p < .*05*, d* = 0.44), within-category for ‘Bike vs. Car’ (Fig.6 green bar: *PER* : *t*(23) = 2.92*, p < .*01*, d* = 0.61), but not for within-category for ‘Male vs. Female’ (Fig.6 orange bar). This pattern of results matches that from the LOC and reflects the structure of perceptual similarity in our stimuli. That is, the most perceptually similar categories (male vs female faces) could not be resolved. Ultimately, these findings are consistent with the idea that the Peripheral ROI encodes low level perceptual stimulus attributes.

**Table 1:**
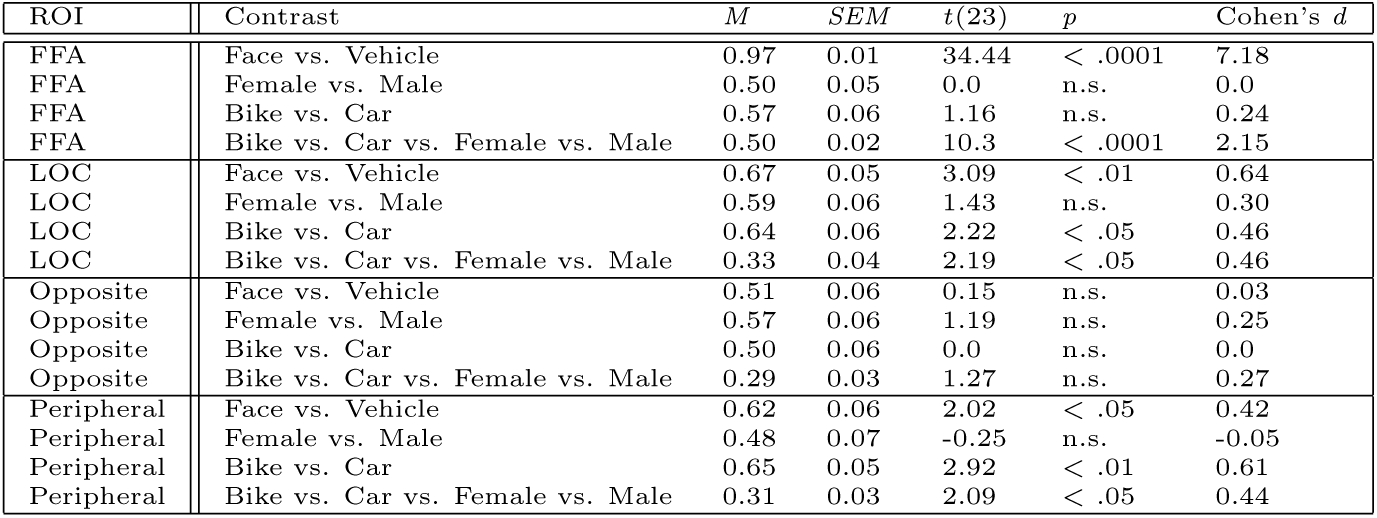
One sample T-test results of the classifier accuracy against chance level (50% or 25%) in each non-Foveal ROI to investigate the classifier ability to discriminate between categories and sub-categories at the group level. M and SEM are, respectively, the average classifier accuracy and the standard error mean for a given contrast.

Fig. 7 displays classifier accuracies across all 8 Foveal ROIs. If information from any particular decoding contrast is present in the fovea, then we expect decoding to be poorest at the smallest ROI, rise to a peak, and potentially fall off at larger ROI sizes depending on the classifier’s success in ignoring irrelevant information. This pattern is most clearly seen for the Face vs. Vehicle contrast (panel A), is also apparent for the Bike vs Car contrast (panel C). It is not present at all for any other contrast (panels B and D). In panels A and C, decoding accuracy reaches a peak at a 2.0° ROI size. As such, we selected the 2.0° fovea ROI to use for comparison to other ROIs in subsequent analyses.

Fig.6 displays classifier accuracies within the 2° Foveal ROI. SVM accuracy was significantly above chance for the category level contrast (‘Face vs. Vehicle’, blue bar: *t*(23) = 1.9*, p < .*05*, d* = 0.4). However, this was not observed for any sub-category or within-category level contrasts (Fig.6. Refer to Table 2). These results clearly establish the presence of feedback from peripheral processing streams to unstimulated foveal areas when viewing real-world stimuli. However, they also suggest that the information relayed from periphery to fovea may be only partial or selective, given certain contrasts decoded in the Peripheral ROI weren’t in the Foveal ROI.

Classification results from the first convolution layer of AlexNet (Fig.4) are qualitatively identical to those obtained in the Peripheral ROI. We found greater than chance SVM accuracy when comparing categories (Fig.4 Face vs. Vehicle, blue bar: *t*(23) = 8.51*, p < .*0001*, d* = 1.2), sub-categories (Fig.4 Bike vs. Car vs. Female vs. Male, red bar: *t*(23) = 5.6*, p < .*0001*, d* = 0.79), within-category vehicles (Fig.4 Bike vs. Car, green bar: *t*(23) = 1.85*, p < .*05*, d* = 0.26), but not for within-category faces (Fig.4 Bike vs. Car vs. Female vs. Male, red bar: *t*(23) = 5.6*, p < .*0001*, d* = 0.79). This result mirrors the patterns of decoding observed in the Peripheral and LOC ROIs. It reinforces the idea that these ROIs are strongly driven by the perceptual features of our stimuli, and further underscores that feedback from peripheral to foveal processing streams may not preserve all information present in the original peripheral stimulus encoding.

**Table 2:** One sample T-test results of the classifier accuracy against chance level (50% or 25%) in each Foveal ROI to investigate the classifier ability to discriminate between categories and sub-categories at the group level. M and SEM are, respectively, the average classifier accuracy and the standard error mean for a given contrast.

| ROI | Contrast | *M* | *SEM* | *t*(23) | *p* | Cohen's *d* |
| --- | --- | --- | --- | --- | --- | --- |
| Foveal 0.5° | Face vs. Vehicle | 0.49 | 0.05 | -0.15 | n.s. | -0.03 |
| Foveal 0.5° | Female vs. Male | 0.49 | 0.06 | -0.14 | n.s. | -0.03 |
| Foveal 0.5° | Bike vs. Car | 0.52 | 0.06 | 0.29 | n.s. | 0.06 |
| Foveal 0.5° | Bike vs. Car vs. Female vs. Male | 0.29 | 0.03 | 1.3 | n.s. | 0.27 |
| Foveal 1° | Face vs. Vehicle | 0.54 | 0.05 | 0.85 | n.s. | 0.18 |
| Foveal 1° | Female vs. Male | 0.44 | 0.06 | -1.01 | n.s. | -0.21 |
| Foveal 1° | Bike vs. Car | 0.57 | 0.06 | 1.16 | n.s. | 0.24 |
| Foveal 1° | Bike vs. Car vs. Female vs. Male | 0.30 | 0.03 | 1.58 | n.s. | 0.33 |
| Foveal 1.5° | Face vs. Vehicle | 0.58 | 0.05 | 1.68 | n.s. | 0.35 |
| Foveal 1.5° | Female vs. Male | 0.42 | 0.06 | -1.35 | n.s. | -0.28 |
| Foveal 1.5° | Bike vs. Car | 0.57 | 0.06 | 1.12 | n.s. | 0.23 |
| Foveal 1.5° | Bike vs. Car vs. Female vs. Male | 0.29 | 0.03 | 1.29 | n.s. | 0.27 |
| Foveal 2° | Face vs. Vehicle | 0.61 | 0.06 | 1.9 | < .05 | 0.4 |
| Foveal 2° | Female vs. Male | 0.43 | 0.05 | -1.28 | n.s. | -0.27 |
| Foveal 2° | Bike vs. Car | 0.58 | 0.06 | 1.29 | n.s. | 0.27 |
| Foveal 2° | Bike vs. Car vs. Female vs. Male | 0.27 | 0.03 | 0.7 | n.s. | 0.15 |
| Foveal 2.5° | Face vs. Vehicle | 0.62 | 0.06 | 2.07 | < .05 | 0.43 |
| Foveal 2.5° | Female vs. Male | 0.47 | 0.05 | -0.66 | n.s. | -0.14 |
| Foveal 2.5° | Bike vs. Car | 0.57 | 0.07 | 1.02 | n.s. | 0.21 |
| Foveal 2.5° | Bike vs. Car vs. Female vs. Male | 0.25 | 0.03 | 0.14 | n.s. | 0.03 |
| Foveal 3° | Face vs. Vehicle | 0.59 | 0.06 | 1.57 | n.s. | 0.33 |
| Foveal 3° | Female vs. Male | 0.46 | 0.05 | -0.85 | n.s. | -0.18 |
| Foveal 3° | Bike vs. Car | 0.58 | 0.06 | 1.29 | n.s. | 0.27 |
| Foveal 3° | Bike vs. Car vs. Female vs. Male | 0.26 | 0.03 | 0.34 | n.s. | 0.07 |
| Foveal 3.5° | Face vs. Vehicle | 0.57 | 0.05 | 1.19 | n.s. | 0.25 |
| Foveal 3.5° | Female vs. Male | 0.48 | 0.04 | -0.37 | n.s. | -0.08 |
| Foveal 3.5° | Bike vs. Car | 0.58 | 0.06 | 1.19 | n.s. | 0.25 |
| Foveal 3.5° | Bike vs. Car vs. Female vs. Male | 0.25 | 0.03 | -0.15 | n.s. | -0.03 |
| Foveal 4° | Face vs. Vehicle | 0.58 | 0.06 | 1.32 | n.s. | 0.28 |
| Foveal 4° | Female vs. Male | 0.50 | 0.05 | 0.0 | n.s. | 0.0 |
| Foveal 4° | Bike vs. Car | 0.55 | 0.06 | 0.78 | n.s. | 0.16 |
| Foveal 4° | Bike vs. Car vs. Female vs. Male | 0.24 | 0.03 | -0.23 | n.s. | -0.05 |

**Figure 6:**
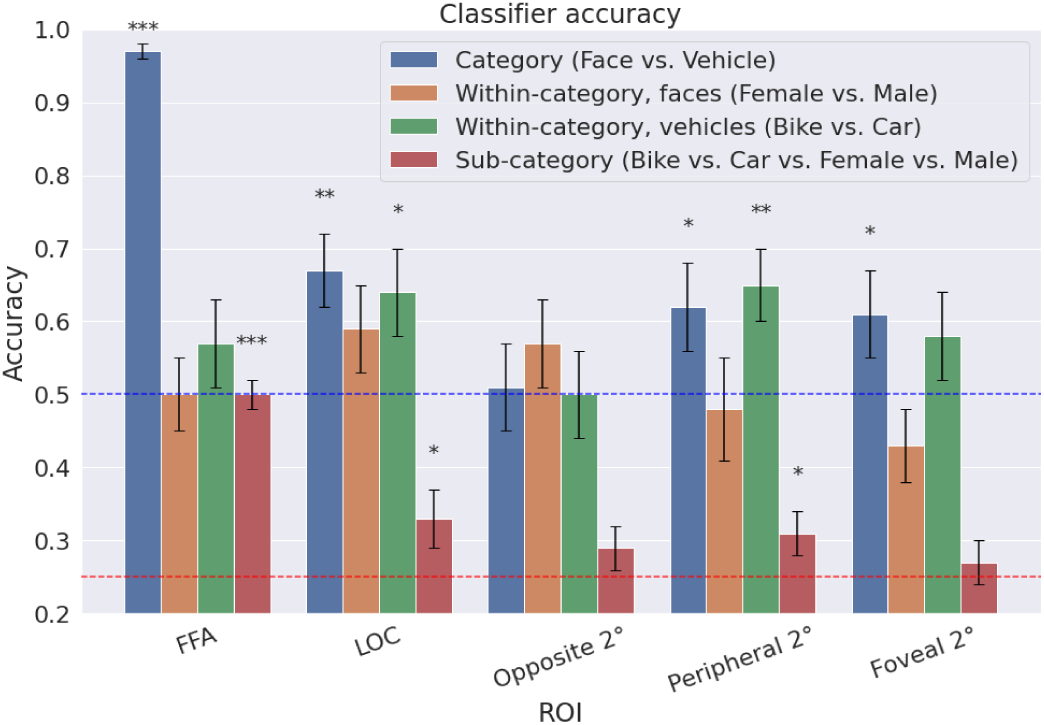
Decoder accuracy for category level (Face vs. Vehicle, in blue), within-category faces ‘Male vs. Female, orange), within category vehicles (Car vs Bike, green), and sub-category level (Bike vs. Car vs. Female vs. Male, in red). Error bars: SEM. Blue line: category level chance (50%). Red line: sub-category level chance (25%). Asterisks (*) indicate statistical significance (e.g., * p < 0.05, ** p < 0.01, *** p < 0.001).

### 3.3. PPI

To examine the functional connectivity between fovea and other visual stream areas, we estimated the mixed effects linear model from Eq. 4 separately for each ROI. The PCA reduced bold time series from the fovea was always used as the target region and the seed ROIs were Opposite, Peripheral, LOC, FFA, and primary auditory cortex (A1). We included A1 as a control condition since it should not be functionally connected to the fovea in our task.

Estimated *β_ppi_* coefficients and corresponding 95% confidence intervals for each ROI are shown in Fig. 8. Inspection of Fig. 8 reveals that *β_ppi_* was significantly greater than zero only in the Peripheral and Opposite ROI, though it was nearly significantly greater than zero in the LOC ROI with the confidence interval for that region narrowly extending over the zero line. The statistical details of these findings are that *β_ppi_* was significantly greater than zero only for the Peripheral ROI (*t*(11.97) = 2.26*, p <* 0.05*, d* = 2.26) and the Opposite ROI (*t*(11.38) = 2.35*, p <* 0.05*, d* = 2.35). It was non-significant for the LOC (*t*(11.60) = 2.002*, p* = 0.07*, d* = 2.00), FFA (*t*(12) = *−*0.65*, p* = 0.53*, d* = *−*0.65), and A1 (*t*(11.71) = *−*0.47*, p* = 0.65*, d* = *−*0.47). The same pattern of results was obtained through parametric bootstrap LRT methods. That is, the full model (including the fixed effect *β_ppi_* term) was preferred over the reduced model (without *β_ppi_*) only in the Peripheral and Opposite ROIs. This was true by all model comparison metrics computed (see Table 3). Thus, we found evidence that fovea was functionally connected to both Peripheral ROIs and did not find evidence that it was functionally connected to any other ROI.

**Figure 7:**
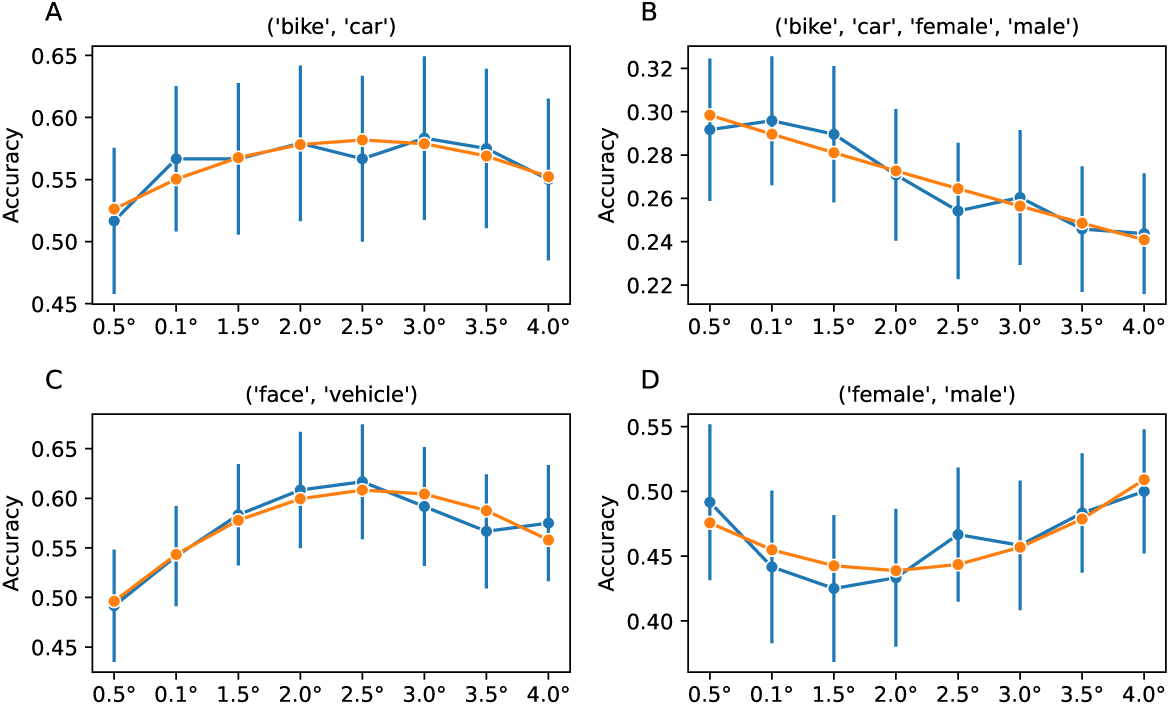
Decoding accuracy across all 8 Foveal ROIs (blue lines) and the best fitting second order polynomial regression line (orange). (A) Results for the within-category vehicles (‘Car’ vs ‘Bike’) contrast. (B) Results for the sub-category level (‘Bike vs. Car vs. Female vs. Male’) contrast. (C) Results for the category level (‘Face vs. Vehicle’) contrast. (D) Results for the within-category faces (‘Male’ vs. ‘Female’) contrast. Error bars indicate SEM.

## 4. Discussion

A growing body of literature demonstrates that information from periphery V1 is projected to foveal V1 and used to aid object discrimination [6, 7, 8, 9, 10, 11]. Here, we hoped to uncover evidence indicating whether this projection is implemented as lateral projections between neurons entirely within primary visual cortex (V1), or is instead implemented by projections from higher-order areas back to lower-order areas. To do so, we explored the role of the foveal cortex in peripheral object discrimination when stimuli were composed of familiar real-world objects, and showed that information about these objects can be decoded in foveal V1, but only if they are characterized by highly dissimilar perceptual representations. This contrasts with peripheral V1, FFA and LOC, all of which provided above-chance decoding of the same stimuli as those decoded in foveal V1, but also permitted decoding of stimuli that where characterized by greater perceptual similarity. Since the pattern of decoding in the fovea was equally dissimilar to that found in lower-order areas (i.e., periphery) as it was to that found in higher-order areas (IT, LOC), these findings are consistent with either periphery-to-fovea neural architecture mentioned above. However, these findings do indicate that feedback from the periphery to the fovea – however it may be implemented – may not preserve all information present in the original peripheral signal. To our knowledge this is the first demonstration of this phenomena. We also performed a functional connectivity analysis that provided evidence that the fovea was functionally connected to both peripheral ROIs (Periphery and Opposite) but not to any other ROI. These results suggest that an implementation contained entirely within V1 may be more plausible.

**Table 3:**
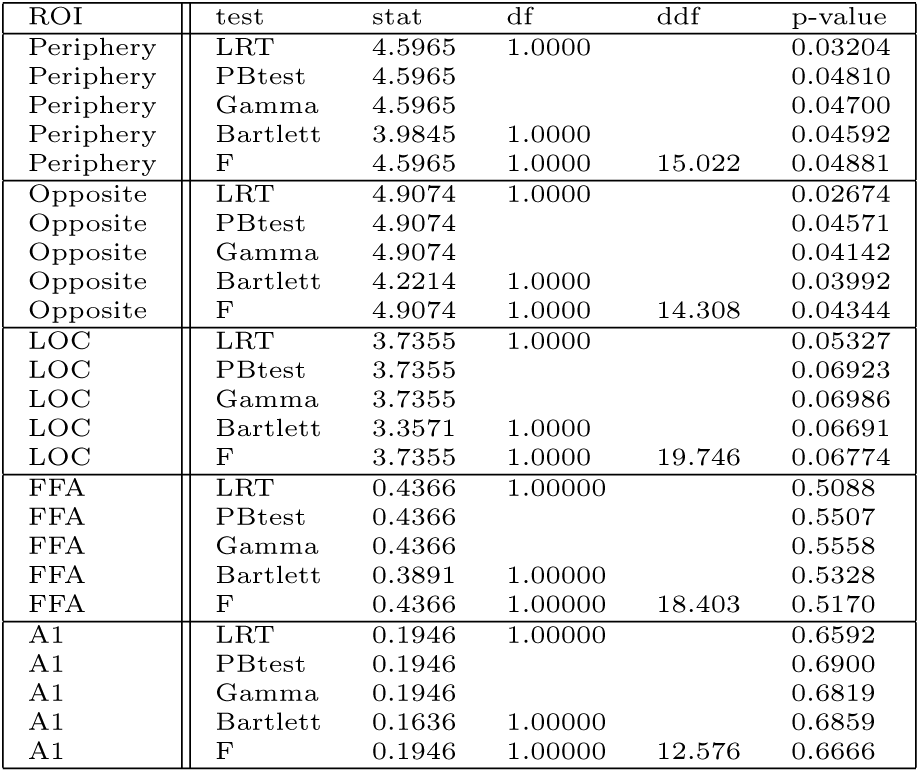
Functional connectivity model comparison results. The models compared are the full model as defined in Eq. 4 and a reduced version of that model with the fixed effect PPI term βppiyppi dropped.

| ROI | test | stat | df | ddf | p-value |
| --- | --- | --- | --- | --- | --- |
| Periphery | LRT | 4.5965 | 1.0000 |  | 0.03204 |
| Periphery | PBtest | 4.5965 |  |  | 0.04810 |
| Periphery | Gamma | 4.5965 |  |  | 0.04700 |
| Periphery | Bartlett | 3.9845 | 1.0000 |  | 0.04592 |
| Periphery | F | 4.5965 | 1.0000 | 15.022 | 0.04881 |
| Opposite | LRT | 4.9074 | 1.0000 |  | 0.02674 |
| Opposite | PBtest | 4.9074 |  |  | 0.04571 |
| Opposite | Gamma | 4.9074 |  |  | 0.04142 |
| Opposite | Bartlett | 4.2214 | 1.0000 |  | 0.03992 |
| Opposite | F | 4.9074 | 1.0000 | 14.308 | 0.04344 |
| LOC | LRT | 3.7355 | 1.0000 |  | 0.05327 |
| LOC | PBtest | 3.7355 |  |  | 0.06923 |
| LOC | Gamma | 3.7355 |  |  | 0.06986 |
| LOC | Bartlett | 3.3571 | 1.0000 |  | 0.06691 |
| LOC | F | 3.7355 | 1.0000 | 19.746 | 0.06774 |
| FFA | LRT | 0.4366 | 1.00000 |  | 0.5088 |
| FFA | PBtest | 0.4366 |  |  | 0.5507 |
| FFA | Gamma | 0.4366 |  |  | 0.5558 |
| FFA | Bartlett | 0.3891 | 1.00000 |  | 0.5328 |
| FFA | F | 0.4366 | 1.00000 | 18.403 | 0.5170 |
| A1 | LRT | 0.1946 | 1.00000 |  | 0.6592 |
| A1 | PBtest | 0.1946 |  |  | 0.6900 |
| A1 | Gamma | 0.1946 |  |  | 0.6819 |
| A1 | Bartlett | 0.1636 | 1.00000 |  | 0.6859 |
| A1 | F | 0.1946 | 1.00000 | 12.576 | 0.6666 |

**Figure 8:**
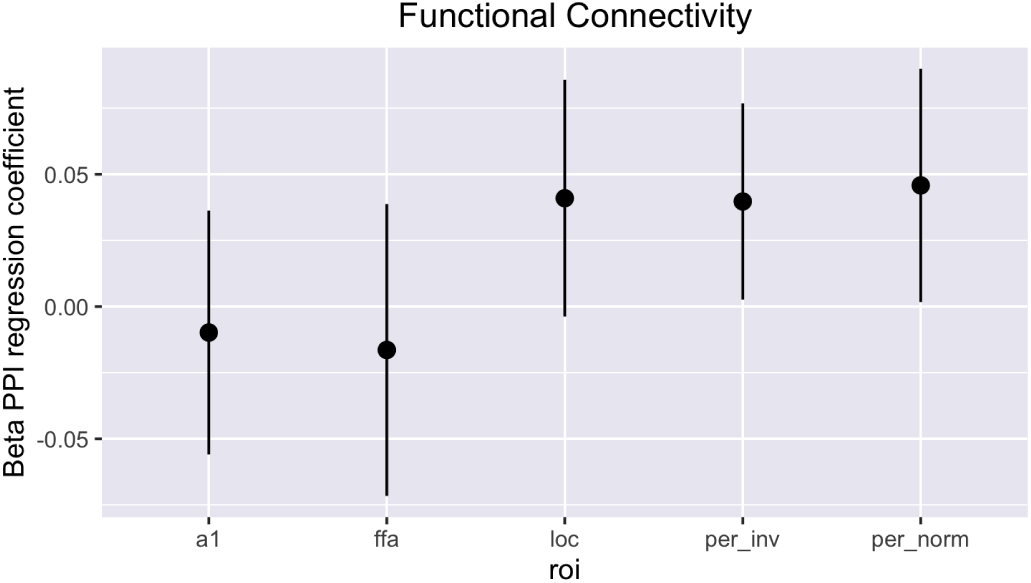
Estimated βppi coefficients and corresponding 95% confidence intervals for each ROI.

### 4.1. Information may be lost along the pathway from periphery to fovea

We were able to decode only whether a given stimulus was a face or a vehicle from the BOLD activity in the fovea ROI. This contrasts with the peripheral ROI (corresponding to where the stimulus was presented in the visual field) for which we were able to decode whether a stimulus was a face or a vehicle, but also whether a vehicle was a bike or a car. This may imply that information is lost along the projection from periphery to fovea. To our knowledge, this is the first report of this phenomena, likely owing to the nature of the stimuli used in our study versus those used in prior studies. Specifically, previous studies all investigated the feedback from periphery to fovea using abstract stimuli that were approximately isomorphic in their perceptual similarities to each other [6, 7, 8, 9, 10, 11]. These stimuli contrast sharply with the real-world stimuli used in this study which produced a considerably more varied distribution of perceptual between-stimulus and between-category similarities (see Fig. 3). Thus, earlier studies were not in a position to detect information loss whereas ours was well suited to this task.

More broadly, our use of real-world stimuli is noteworthy on its own merits. In particular, that we can decode anything at all about real-world images clearly demonstrate that the presence of peripheral information in the foveal cortex is not dependent upon experimental paradigms using abstract unfamiliar stimuli. It also indicates that the familiar categories intrinsic to real-world stimuli do not seem to trigger neural processing that interferes with or negates feedback from periphery to fovea. This finding lends substantial credence to the idea that peripheral projections to foveal cortex may be at play in real-world behavior.

### 4.2. Neural architecture of periphery-to-fovea feedback

We sought to resolve whether feedback from periphery to fovea visual areas is implemented by local circuits within V1 versus as projections from higher-order areas back to lower-order areas. We took a two-pronged approach to meet this goal. First, we compared the pattern of decoding observed in the fovea V1 to those obtained in periphery V1, FFA, and LOC. Similarity to the decoding pattern obtained in periphery V1 would be aligned with a local V1 circuit implementation whereas similarity to the decoding pattern obtained in FFA or LOC would be aligned with a higher area to lower area implementation. We observed a pattern of decoding in the fovea that was equally dissimilar to that obtained in all other visual regions explored. As such, the pattern of decoding observed here does not provide evidence that is more consistent with either neural architecture.

However, one feature limiting our ability to make the above comparisons with confidence is that we were not able to decode male faces from female faces in any ROI. Given the high perceptual similarity of these stimuli, it is not surprising that we could not decode them in peripheral V1. It is, however, surprising that we could not decode them even in FFA [58]. Since our imaging protocol and analysis methods are not very different from prior studies in which such decoding was observed, we suspect that our inability to decode male versus female faces in FFA most likely reflects the short presentation time and the peripheral location of the stimuli. For example, there may be impoverished encoding of the visual stimuli in V1 due to the peripheral neurons’ lower acuity and the fast presentation time of the stimuli. Furthermore, evidence suggest that face perception, and specifically gender and identity perception, might arise as a result of a more complex network that, along with FFA, include – but it is not limited to – brain regions not considered in our analysis, such as the Occipital Face Area [59, 60]. Finally, neurons within face-related networks have been found to exhibit a foveal bias, as opposed to peripheral bias seen in scene perception neurons. This preference for centrally presented stimuli could in turn impact the processing of perceptual and categorical features in FFA for our peripherally presented stimuli [61]. Future studies should explore these possibilities.

We also assessed the neural architecture of the feedback from periphery to fovea by performing a PPI functional connectivity analysis. This analysis revealed that the fovea was significantly functionally connected only to the periphery. It was not significantly functionally connected to any other visual ROI considered, nor was it functionally connected to primary auditory cortex which we included as a control. These results favour a local circuit implementation of periphery-to-fovea feedback. Furthermore, an interesting aspect of this result is that fovea was significantly functionally connected to both perihperal ROIs (Perihperal and Opposite). It is possible to imagine any number of network architectures that may produce such a pattern of results and we hope to direct future research at this issue.

Finally, another aspect of our results may also be more consistent with the local V1 circuit implementation. In particular, our results show that information from peripheral retinotopic areas is fed back into the foveal cortex (or, at least, can be decoded in the foveal cortex) only in an anatomical region corresponding to the central 2-2.5 visual degrees of the visual field. Interestingly, this size approximately matches the average size of the peripheral stimuli presented during the behavioral task (1.8 visual degrees, see section 2.8). This striking similarity suggests that the information fed back to the foveal cortex resembles information shaped by the receptive field properties of peripheral V1 circuits more than those of higher order visual areas.

### 4.3. Implications for current models and theories

Traditional feedforward views of visual object recognition posit that visual information is passed along a processing stream of sequential non-linear transformations from lower to higher visual areas beginning in the retina and ending in higher-order cognitive areas [62, 63, 64]. However, feedback and recurrence are central design features of the human brain, and are heavily expressed in the human visual system. Almost since their discovery, researchers have focused on understanding the role of these extensive feedback connections from higher to lower cognitive and visual areas (e.g., attentional models [65, 66, 67] and predictive coding models [68]).

Feedback models of vision posit a modulatory role of feedback connections over the perceptual (bottom-up) input. For instance, attentional models theorise a central role of feedback connection in selective attention [66] over task-relevant features – and therefore a facilitation based on internal goals. On the other hand, predictive models theorise a top-down modulatory effect of feedback connections based on prior knowledge of the world. In this context, core object recognition would still be carried by the feedforward sweep, while high-level areas make predictions (i.e., hypothesis) about the visual scene to modulate activity in low-level areas [69, 70].

However, recent evidence – including the results of the present study – suggests that the idea of restricting feedback connections to a modulatory role of the visual input might be too limited. For instance, Williams et al. [6] found information about peripherally presented stimuli in the (unstimulated) foveal region of the primary visual cortex. This new phenomenon has been corroborated by several TMS [7] and behavioral experiments [9, 10, 11]. These authors reported a new type of feedback, not predicted by previous models, that does not modulate the perceptual information but rather creates a completely new task-relevant and position-invariant representation in a brain area with no direct visual input.

Although the precise nature of periphery-to-fovea feedback is unclear, several frameworks have been proposed to explain top-down effects in V1. On the one hand, they might reflect pre-saccadic mechanisms, where behaviorally relevant aspects of the peripheral pre-saccadic information are fed back to the foveal cortex before the saccade, and integrated with the post-saccadic foveal stimulus [71, 72, 73]. On the other hand, they might reflect a predictive generative mechanism, where high-level areas generate a perceptual hypothesis of the peripheral stimuli based on prior knowledge, and feed this information back to the foveal V1 to facilitate representations consistent with the selected hypothesis (i.e., to re-weights bottom-up visual representations) [74, 75, 76, 77, 78, 79, 71, 72].

An appealing aspect of the generative hypothesis is that the process of prediction may result in a compression of the information from high-order areas to lower ones that may strip the original representation from some perceptual features in order to feed back only task-relevant information as a pre-saccadic modulatory mechanism. This account therefore seems consistent with the decoding differences we observed between the peripheral ROI, where the activation pattern primarily stems from the encoding of the stimulus, and the foveal ROI, where it could be a result of a generative process prone to information loss. For instance, this feedback might represent a synthetic summary of the visual stimuli, where only *some* of the original perceptual features are represented (e.g., shape, color, luminance/contrast, low spatial frequency, etc.). This compression can, in principle, affect decoding analyses, and it is in line with our results showing both significant decoding for within-category vehicles classifications (cars vs. bikes) in the peripheral ROI but not in the foveal cortex and with previous work showing task-dependent information in the foveal cortex during similar tasks [6, 9, 10]. However, it is difficult to reconcile the idea that this generative process is driven from higher-level visual areas (i.e., that it is top-down) with our observation of functional connectivity only between ROIs within primary visual cortex.

Finally, Lee et al. [69] have proposed a Bayesian framework where each area is specialized in processing some aspects of the stimulus, with V1 being a “high resolution buffer”. Each area generates *posteriors* to influence bottom-up *priors* in order to constrain the possible space of a perceptual hypothesis through a reciprocal interaction. Considering the high-resolution specialization of V1, and the continuous reciprocal interactions between priors and posteriors, this framework predicts high-resolution perceptual information carried by the periphery-to-fovea feedback, and may therefore be difficult to reconcile with our observation that information appears to be lost somewhere along its path from periphery to fovea.

### 4.4. Limitations and future directions

One limitation of the present work is that a low signal-to-noise ratio, common in unprocessed or minimally processed fMRI data, can lead to spurious patterns of results. However, we have no reasons to think that noise components in the data may drive our pattern of results, and possible strategies to mitigate this issue (such as heavier data denoising pipelines) could, on the other hand, have a detrimental impact on the fMRI activation patterns and, therefore, on the multi-voxel pattern analysis performed.

A second limitation of the present work is that the observed pattern of results may be a consequence of the classifier failing to detect existing differences. Specifically, we cannot conclude with absolute certainty that the comparisons that the classifier is failing to distinguish with an accuracy higher than chance (e.g., male vs. female faces in the foveal cortex) are, in fact, indistinguishable in the brain. These conditions may, in principle, be perfectly classifiable using different neuroimaging methods with a higher temporal or spatial definition or, on the other hand, Support Vector Machine algorithms may just lack the sensitivity needed to pick up such existing differences due to the intrinsic neural population coding for specific categories or features. However, this may be unlikely given other studies have successfully decoded information about male vs. female faces in several brain regions [58].

Signal-to-noise ratio concerns are related to overall statistical power concerns. One area where this seems particularly relevant is in the results of the PPI functional connectivity analysis. In particular, the peripheral ROIs are just barely significant while the LOC is just barely non-significant (see Fig. 8). However, all three of these areas appear qualitatively distinct from the FFA and A1 ROIs. Thus, it is possible that the LOC was functionally connected to the fovea in the present data, but we simply lacked the power to detect it.

Finally, the present work leaves several important questions to be addressed by future research. Foremost among these might be to perform an-other neuroimaging study in which not just the perceptual similarities of stimuli to each other is manipulated, but their relevance to task performance is also manipulated. This should produce deeper insight into the nature of information loss in periphery-to-fovea feedback. Another important avenue for future research may be to acquire fMRI data from imaging protocols optimized to detect functional connectivity and causality in visual circuits. Doing so may shed greater light and confer greater confidence in the evidence for or against the idea that our data seems to suggest that periphery-to-fovea feedback is implemented within circuits local to V1.

## 5. Data and Code Availability

All data necessary to reproduce the primary findings and plots presented in this study are available for public download. The (minimal) dataset, which has been structured following BIDS naming convention, can be accessed at https://osf.io/h95a2/. The code underpinning the analyses and results of this study has been made available to the community, and it can be retrieved from the following repository: .

## 6. Author Contributions

AC: Conceptualization, Data curation, Formal analysis, Investigation, Methodology, Project administration, Resources, Software, Validation, Visualization, Writing – original draft, and Writing – review & editing. MW: Conceptualization, Methodology, Project administration, Supervision, and Writing – original draft. MC: Formal analysis, Methodology, Project administration, Software, Supervision, Validation, and Writing – original draft, review & editing. BT: Formal analysis, Methodology, and Writing – review & editing.

## 7. Funding

This research did not receive any specific grant from funding agencies in the public, commercial, or not-for-profit sectors.

## 8. Declaration of Competing Interests

The authors declare no competing interests.

## 9. Supplementary Material

**Supplementary Figure 1:**
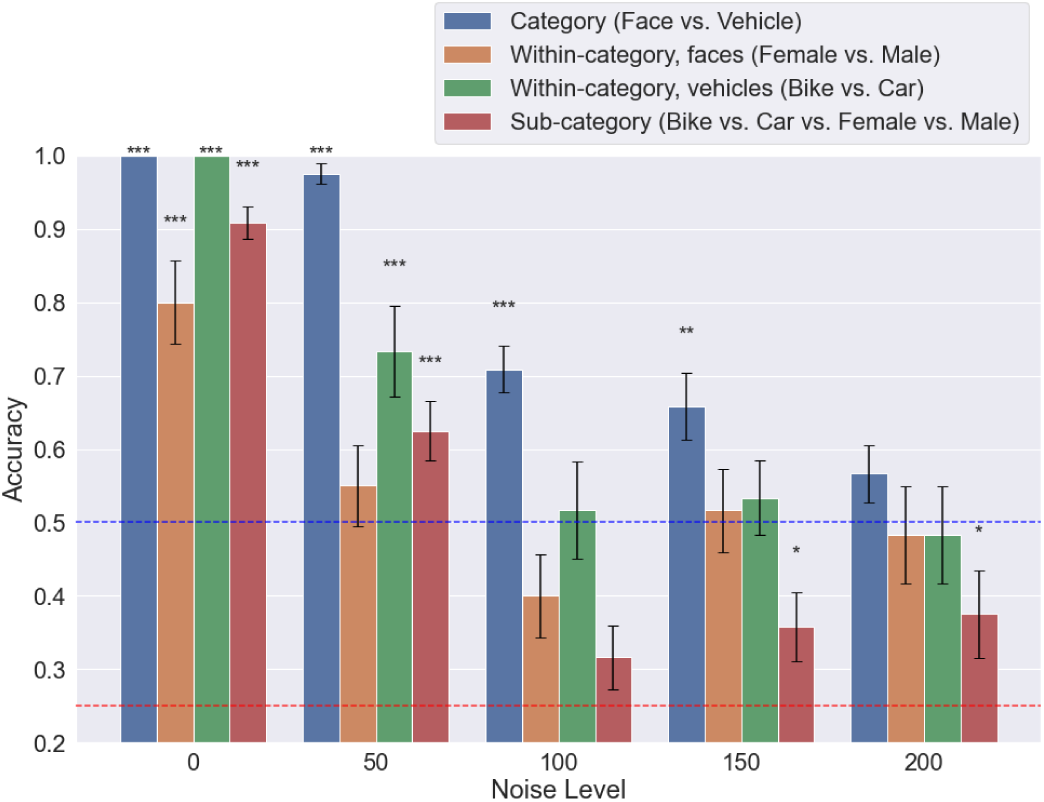
Decoder accuracy in the perceptual model at different noise sigmas for category level (’Face vs. Vehicle’, in blue), within-category faces (‘Male’ vs. ‘Female’, orange), within category vehicles (‘Car’ vs ‘Bike’, green), and sub-category level (‘Bike vs. Car vs. Female vs. Male‘, in red). Error bars: standard error mean (SEM). Blue line: category level chance (50%). Red line: sub-category level chance (25%).

**Supplementary Figure 2:**
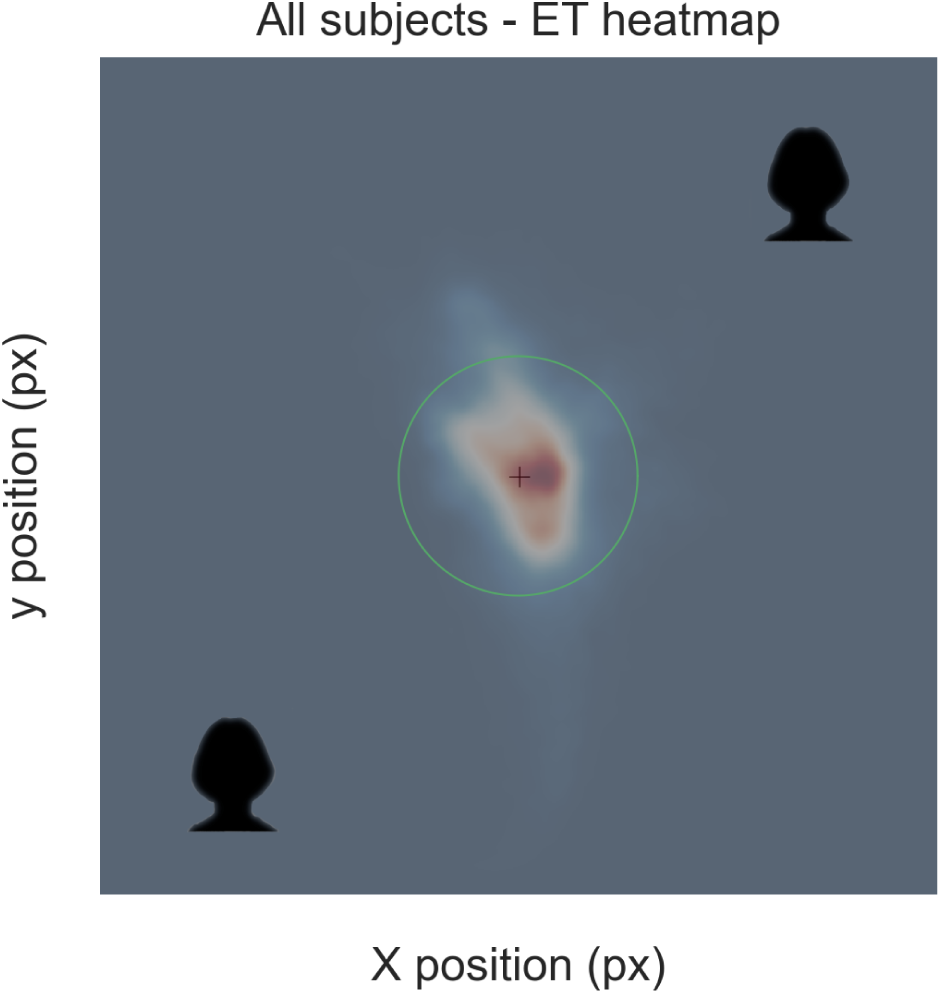
Faces are obscured due to the journal policy. Average fixation heat-map during the fMRI experimental task across all subjects and runs. The green circle represents the area within 2 visual degrees radius from the central fixation point.

## Footnotes

1 Although a classifier might learn to ignore the irrelevant dimensions, it’s still beneficial to minimize irrelevant information in decoding applications.

## Notes

https://github.com/costantinoai/foveal-feedback-2023

### Competing Interest Statement

The authors have declared no competing interest.

